# Targeted protein degradation of the CPSF complex by benzoxaboroles through sumoylation

**DOI:** 10.1101/2024.03.11.584402

**Authors:** Ning Zhang, Abdul Samad, Farnaz Zahedifard, Richard J. Wall, David Horn, Martin Zoltner, Mark C. Field

**Affiliations:** School of Life Sciences, University of Dundee, Dundee, DD1 5EH, UK; Department of Parasitology, Faculty of Science, Charles University in Prague, BIOCEV, Vestec, Czech Republic; Biology Centre, Czech Academy of Sciences, Institute of Parasitology, České Budějovice, Czech Republic

**Keywords:** SUMO, CPSF complex, benzoxaborole, proteasome, targeted protein degradation, mode of action, trypanosoma

## Abstract

Benzoxaboroles (BoBs) feature a boron-heterocyclic core and are an important innovation in the development of drugs against a range of pathogens and other pathologies. A broad spectrum of pharmacology is associated with chemically diverse BoB derivatives and includes multiple modes-of-action (MoA) and targets. However, a consensus MoA for BoBs targeting evolutionarily diverse protozoan pathogens has emerged with the identification of CPSF3/CPSF73 in the CPSF complex in both apicomplexan and kinetoplastida parasites. Here we establish a functional connection between protein sumoylation and the boron-heterocyclic scaffold shared by all BoBs using comprehensive genetic screens in *Trypanosoma brucei.* There is a rapid temporal and spatial shift in global protein sumoylation following BoB exposure and members of the CPSF complex are specifically destabilised in a SUMO and proteosome-dependent manner. Finally, we find rapid decrease in bulk mRNA levels, consistent with the role of CPSF3 in mRNA maturation. We propose that a combination of direct inhibition coupled with targeted degradation of CPSF3 underpins the specificity of BoBs against trypanosomatids.

## Introduction

Current and emerging pathogens threaten public health, and neglected tropical diseases (NTDs) remain a significant concern to much of the world. Collective efforts have decreased the global burden of several NTDs, including African trypanosomiasis and malaria, with the former at historical lows. Cases of both diseases have fallen considerably in the past decade, *albeit* with concerns that pressures from COVID-19 and additional geopolitical events will facilitate a resurgence (Jamabo et al., 2023, Solano et al., 2023).

Two new compounds recently entered the pipeline for the treatment of African trypanosomiasis; fexinidazole and acoziborole (Solana et al., 2023). Acoziborole is a benzoxaborole (BoB), a class that recently emerged as potent microbicides against a wide spectrum of infections, spanning viral, bacterial, protozoan and fungal agents (Nguyen et al., 2023, Tarleton 2023, Huang et al., 2023). Significantly, BoBs demonstrate activity towards both kinetoplastida and apicomplexan parasites, despite their wide evolutionary divergence. A common target, cleavage and polyadenylation specificity factor 3 (CPSF73/CPSF3/YSH1) has been identified for BoB activity against multiple protists (Sonoiki et al., 2017, Palencia et al., 2017, Wall et al., 2018, Mowbray et al., 2021). CPSF3 is widely conserved, with endonuclease activity recognising the pre-mRNA 3’-cleavage site AAUAAA prior to polyadenylation (reviewed in Swale and Hakimi, 2023). Amongst many identified functions, CPSF3 also possesses 5’ to 3’ exonuclease activity and operates in a U7 snRNP-dependent manner. Moreover, in higher eukaryotes such as *Arabidopsis thaliana,* CPSF3 also participates in microRNA maturation (Has et al., 2023). The full range of CPSF complex functions remain to be determined, and our understanding of CFPS3 activity in protists is rudimentary, complicated further by the high degree of cooperation between proteins that process RNAs for export to the cytoplasm, making the roles of individual proteins difficult to discern (Inoue et al., 2022).

CPSF3 is the sole catalytic component of the CPSF complex and the target of several distinct BoBs in multiple unicellular protozoan parasites, including *Plasmodium falciparum, Toxoplasma gondii, Trypanosoma brucei* and *T. cruzi* (reviewed in Swale and Hakimi 2023, Has et al., 2023). Evidence that CPSF3 represents the trypanosome target is supported by demonstrations that *trans*-splicing is inhibited by multiple BoBs (Begolo et al., 2018, Waithaka and Clayton 2022) and from overexpression screens and drug resistance-associated mutation at the drug-binding site (Wall et al., 2018). Significantly, CPSF3 is not the target for BoB action in bacteria or fungi, *albeit* that this most likely depends on additional determinants in the BoB structure as well as biochemical factors (Ganapathay et al., 2023). Several metabolic pathways in trypanosomes process and activate specific BoB derivatives, serving to retain them within the parasite and hence potentiate trypanocidal activity (Zhang et al., 2018; Giordani et al., 2020). BoBs also impact S-adenosyl-L-methionine metabolism, and hence likely many methylation events (Steketee et al., 2018)

A number of features distinguish transcriptional mechanisms in kinetoplastids from animals, plants and fungi. In particular polycistronic transcription and *trans*-splicing shifts the major mRNA regulatory functions from *cis*-acting promoter elements onto post-transcriptional steps including processing, maturation, quality control and export as well as *cis*-based signals within UTRs that mediate turnover and translatability (e.g. Zoltner et al., 2018, Goos et al., 2019). Unlike animals and fungi, inhibition of splicing leads to a pool of mRNAs, in the form of granules located proximal to the cytoplasmic side of the nuclear pore complex (NPC), in contrast to retention within the nucleus (Goos et al., 2019). The trypanosome export machinery associated with the NPC lacks the DBP5 RNA helicase Gle1 and other cytoplasmic NPC components that constitute the mRNA quality control (QC) platform in animals and fungi, suggesting that these cytoplasmic mRNA foci represent an analog of the canonical NPC-associated QC system (Obado et al., 2022). There are multiple pathways mediating processing of distinct RNA cohorts in trypanosomes. For example, two distinct polyA-binding protein complexes share few components (Zoltner et al., 2018) while specificity within interactomes identified for trypanosome eIF4III, Sub2 and additional complex components has been demonstrated (Inoue et al., 2022). Many canonical factors are absent and likely replaced by lineage-specific proteins.

SUMO (small ubiquitin-like modifier) is widely distributed across eukaryotes and influences protein location, stability and function (Flotho and Melchior 2013). In most taxa, including trypanosomes, SUMO is located primarily in the nucleus. Humans possess four SUMO paralogs but trypanosomes have only a single gene for SUMO as well as the E1 and E2 SUMO ligases. In *T. brucei* SUMO is closely associated with transcriptional and post-transcriptional regulation proximal to the expression site body (ESB) (Iribarren et al., 2018, Saura et al., 2019) where transcription of the superabundant variant surface glycoprotein (VSG) takes place and represents a major mRNA processing site, together with production of the spliced leader (López-Farfán et al., 2014, Faria et al., 2021). Significantly, the ESB contains multiple proteins responsible for securing monoallelic expression of VSG, while sumoylation is associated with proteins located at VSG-encoding chromatin (Saura et al., 2019). SUMO is important for translocation of mRNA to the NPC in animals and fungal cells (Gasser and Stutz 2023), and while not demonstrated experimentally, there is sufficient conservation within trypanosomes to expect that similar mechanisms operate.

Generation of resistant mutants and genome-wide screens have identified two classes of trypanosome genes implicated in BoB sensitivity. These prodrug processing mechanisms are mediated by parasite aldehyde dehydrogenase (Zhang et al., 2018) or serine carboxypeptidase (Giordani et al., 2020), and are sensitive to BoB structure. Here we describe a further cohort that are essentially insensitive to structure beyond the BoB core and are associated with the sumoylation pathway. Furthermore, BoB exposure triggers rapid hypersumoylation, mediating depletion of some CPSF complex components, including CPSF3. Further, sumoylation plays an essential role in regulating the subcellular localisation of the CPSF complex. This dual impact on CPSF3, mediated by a direct inhibition of enzyme activity and targeted degradation, likely accounts for benzoxaborole trypanocidal potency.

## Results

### The protein sumoylation pathway underpins sensitivity to benzoxaboroles

A frequently identified BoB target is CPSF3, and which significantly has been indicated for a diverse range of protist pathogens. This evidence suggests a conserved mechanism underpinning BoB toxicity (Sonoiki et al., 2017, Palencia et al., 2017, Wall et al., 2018, Mowbray et al., 2021). Evidence is also consistent with the direct inhibition of CPSF3 function, based on *in silico* modeling, mutagenesis and phenotypic data that indicates compromised *trans*-splicing in trypanosomes, and also confirming that CPSF3 acts within mRNA maturation in trypanosomes (Begolo et al., 2018, Waithaka and Clayton 2022). However, a more complex interaction between BoBs and trypanosomes may be relevant and specifically the modest level of resistance obtained by overexpression of CPSF3, an approximate threefold change to EC_50_, compared with more than a hundredfold for a block to a prodrug processing pathway suggests a possible additional factor is important (Zhang et al., 2018, Wall et al., 2018). To probe this possibility, we systematically explored structure-activity relationships (SAR) for a range of benzoxaboroles by challenging the African trypanosome bloodstage form. Clinically-relevant together with structurally diverse BoBs were selected for an unbiased genome-wide RNAi (RIT-seq) screen.

A common cohort of genes was identified in the majority of screens, *albeit* not all to the same magnitude. These include TbSUMO (Tb927.5.3210), TbUBA2/E1 (Tb927.5.3430) and TbUBC9/E2 (Tb927.2.2460) (Figure 1, highlighted in red) (Ye et al., 2015). A fourth gene (Tb927.11.6340) was also identified by AN6306 and AN9872. This gene encodes a protein predicted to contain two peptide N-glycanase/UBA or UBX-containing protein domains (PUB), which we designate as d(ouble)PUBP and is orthologous to UBXN7/UBXD1, a cofactor of the AAA-ATPase VCP/p97/Cdc48 and possessing additional functions within the ubiquitylation pathway (Tarcan et al., 2022, Di Gregorio et al., 2021). Significantly, dPUBP was also identified as a gene sensitizing trypanosomes to apolipoprotein-L1/TLF, a key host restriction factor for African trypanosome infection (Alsford et al., 2014). AN10070 and AN11736 had comparatively low magnitude for SUMO pathway genes; significantly both compounds have a methyl substituent at position 7 on the benzoyl ring of the core BoB pharmacophore, which may impact their interaction with CPSF3.

**Figure 1:**
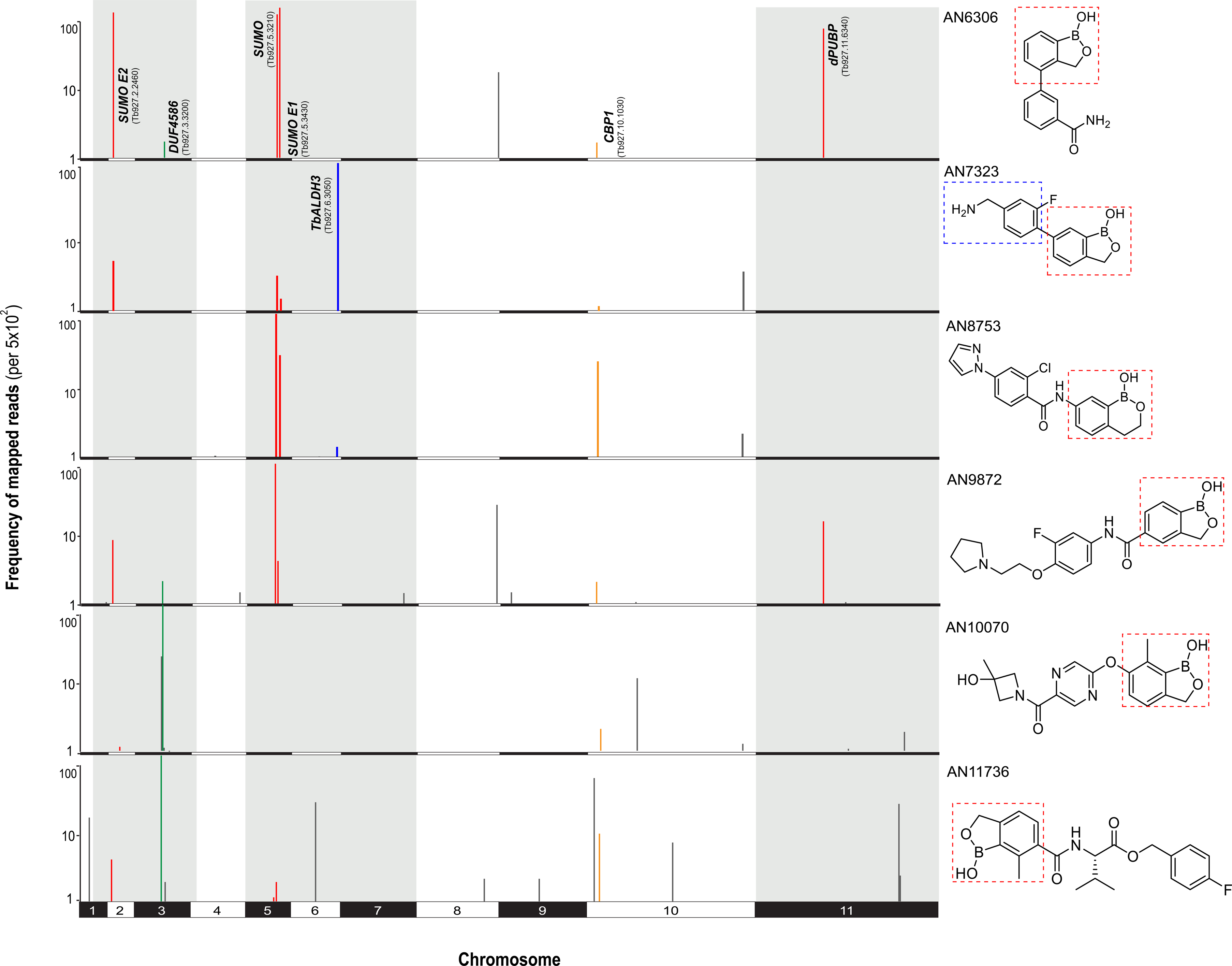
A whole genome RNAi screen identifies genes associated with benzoxaborole MoA. Manhattan plots for genes linked to benzoxaborole sensitivity identified across the trypanosome genome. Populations that survived drug exposure were collected for barcode sequencing. Plots indicate frequency of mapped reads against their positions in the genome, represented for the eleven megabase chromosomes. Data are filtered through a designated cut-off of >500 RPM and must be identified with >2 independent knockdown sequences to be registered as a significant hit. Red indicates all genes identified related to protein sumoylation, the blue line indicates a gene specific to AN7323, green are genes predominantly recovered for AN10070 and AN11736, while yellow indicates CBP1. At right are the chemical structures of the compounds included in this screen. The heterocyclic benzoxaborole scaffold is indicated with the red box and the benzoylmethylamine moiety of AN7323 is highlighted with a blue box. Full RITseq read data are available in Table S1.

Notably, AN7323, carrying a methylamine substitution, also identified TbALDH3, which is involved in prodrug activation for AN3054 in combination with a host-encoded monoamine oxidase (Zhang et al., 2018). Further, AN6306, AN8753, AN9872, AN10070 and AN11736 all identified CBP1 (Tb927.10.1030) which acts on peptide bonds and is a further prodrug activation pathway (Giordani et al., 2020). The low number of CBP1 reads for AN6306 and AN10070 may indicate poor CBP1 activity against non-peptidic amine-containing structures. For AN9872, AN10070 and AN11736 additional resistance determinants were identified on chromosome 3 (Figure 1 green). These latter are not considered further here.

We analysed an additional cohort of BoBs by RIT-seq to extend our dataset; these include the previously characterised TbALDH3 isomeric substrates AN3054 and AN3057 together with acoziborole (AN5568) (Figure 2A, Table S1). All three compounds also identified SUMO pathway genes (red) whilst AN3054 and AN3057, as expected, identified TbALDH3 (blue) as well as Tb927.11.8870, a mitochondrial DEAD-box protein (blue). Taken together analysis of this cohort of BoBs indicates a common role for SUMO pathway proteins in seven out of nine screens, with the two compounds lacking evidence for SUMO interactions being characterised by a methyl substituent on the BoB core.

**Figure 2:**
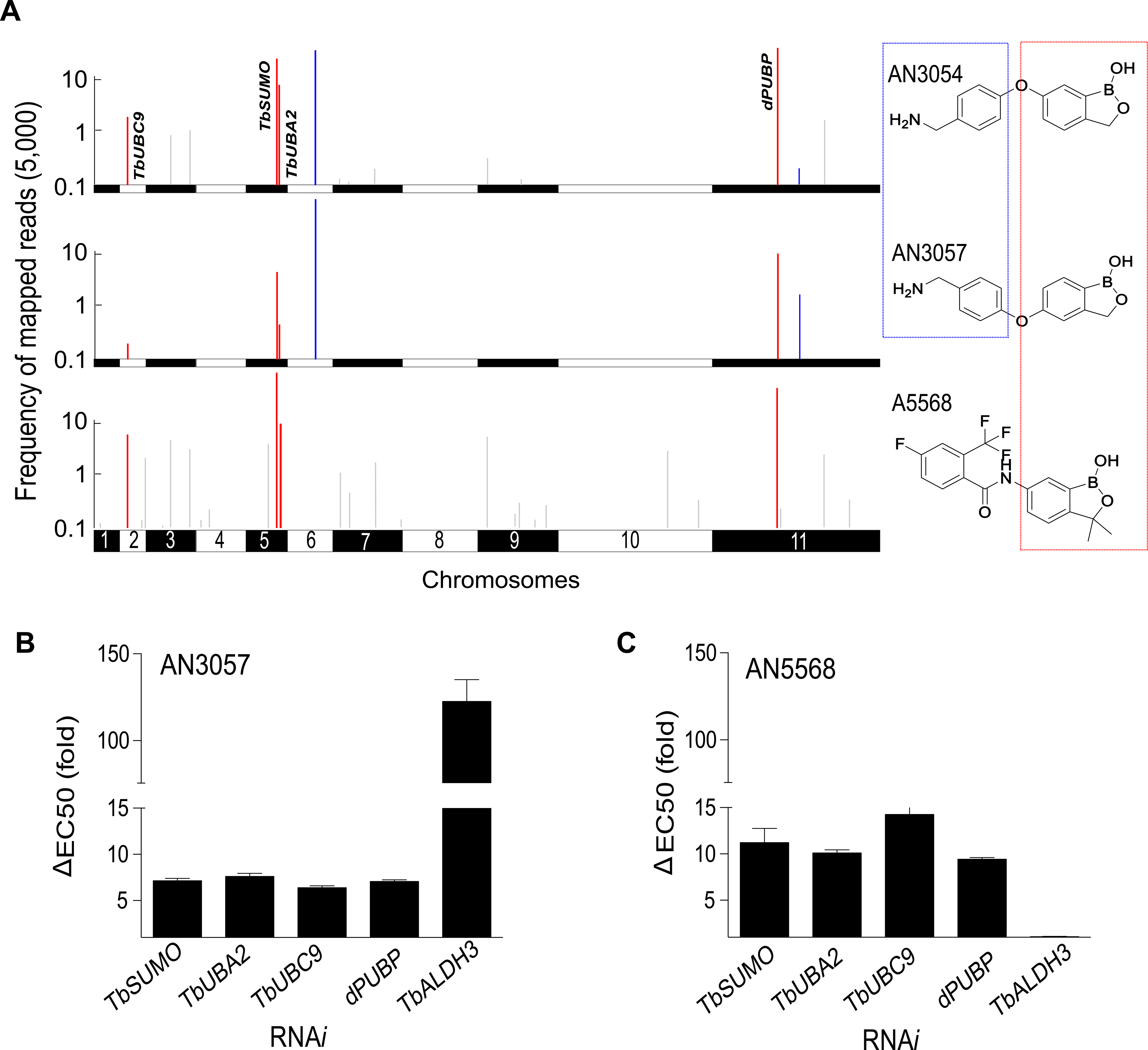
Additional RNAi screen confirms genes associated with benzoxaborole MoA. A: Manhattan plots of genes linked to benzoxaborole sensitivity identified across the trypanosome genome, processed and presented as described for Figure 1. Red indicates genes related to protein sumoylation and blue indicates hits specific to AN3057 and AN3054. At right are the chemical structures of the compounds included in this screen. The heterocyclic benzoxaborole scaffold is indicated with a red box and the benzoylmethylamine moiety is highlighted with a blue box. Full RITseq read data are available in Table S2. B. Resistance to the benzoxaboroles in parasites with silenced protein sumoylation factors. The changes in EC_50_ of the designated compound was measured for cells prior to and post induction of dsRNA.

To validate that the SUMO pathway is an active determinant for BoB sensitivity we used RNAi to specifically target the individual genes identified. This demonstrated that both AN3057 and acoziborole required SUMO pathway gene products, including dPUB, for full sensitivity to BoB trypanocidal activity, with an approximate five and ten-fold alteration to the EC_50_ respectively (Figure 2B, Figure S1). As expected, only AN3057 was impacted by TbALDH3 silencing. Collectively, these gene products constitute a near complete SUMO-conjugation pathway, specifically SUMO and E1 activating and E2 conjugating factors. The absence of TbSiz1 (Tb927.9.11070), the only characterised SUMO E3 ligase of trypanosomes (López-Farfán et al., 2014) from all of our screens suggests that the mechanism here may rely on direct E2-conjugation to substrates (Lin et al., 2004).

### Benzoxaboroles elicit a hypersumoylation response

We analysed the levels of sumoylated/ubiquitylated proteins using Western blotting of whole cell extracts with antibodies to both SUMO and ubiquitin, following exposure to BoB, silencing of the SUMO E1, E2 or dPUBP, or both treatments. In the absence of any treatment, a spectrum of sumoylated proteins were detected, with molecular weights exceeding ∼250 kDa and these sumoylated proteins were globally decreased upon knockdown of the SUMO E1, E2 or dPUBP (Figure 3A). This clearly indicates that the genes identified in the screens indeed encode components of the sumoylation pathway. Significantly, for E1 or dPUBP knockdown, BoB treatment effectively protected against decreased sumoylation, but not for the E2 knockdown (Figure 3A).

**Figure 3:**
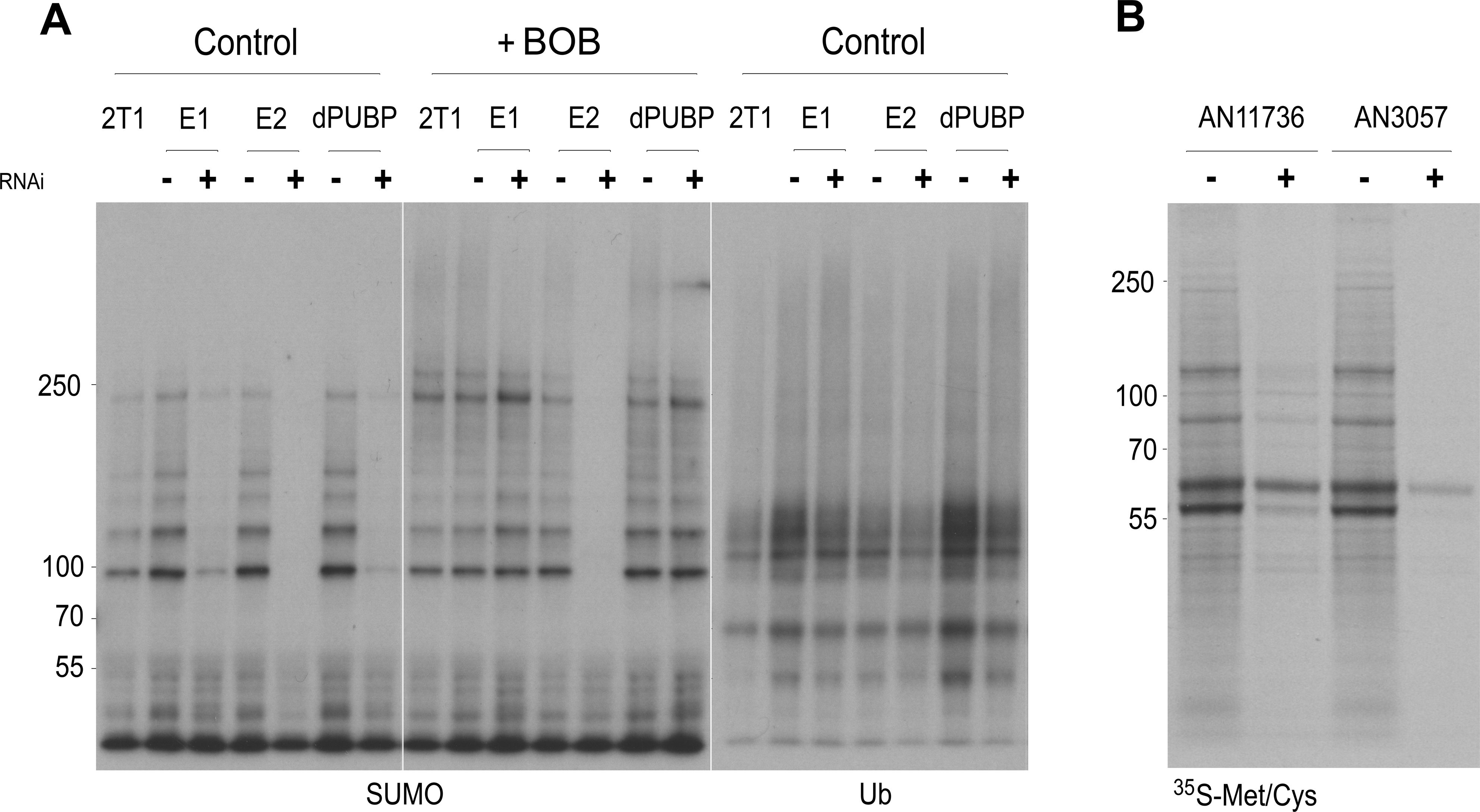
Benzoxaborole exposure leads to inhibition of protein synthesis. A: Western blot of whole cell lysates exposed to benzoxaboroles and probed with anti-SUMO antibody. Cells with and without silencing for SUMO pathway components identified from RITseq screens were analysed. Note that on the left the SUMO adducts are significantly reduced by knockdown, but in the presence of benzoxaborole the SUMO levels are not significantly altered for E1 or dPUBP. B: Duplicate lysates probed with anti-ubiquitin antibody. Ubiquitylation remains present in silenced cells, albeit with a decrease in intensity, likely a result of reduced protein synthesis. C: ^35^SMet/Cys metabolic-labeling in the presence and absence of indicated benzoxaboroles, demonstrating reduced incorporation of radiolabeled amino acids in compound treated cells. The major bands at 60kDa and 65kDa likely correspond to tubulin and VSG respectively. At left are molecular weight markers, and 2T1 refers to the parental cell line without an RNAi construct.

Probing of identical lysates with anti-ubiquitin antibodies demonstrated that the impact was essentially SUMO-specific, as there were only modest impacts to levels of ubiquitylated proteins upon silencing (Figure 3A). A decrease in the ubiquitin signal is likely a result of decreased protein synthesis, as demonstrated by the significant loss of incorporation of ^35^S-Met/Cys into protein on BoB exposure (Figure 3B), indicating a collapse in protein synthesis and consistent with inhibition of *trans*-splicing of protein coding transcripts.

### Benzoxaboroles impact CPSF3 stability

We monitored dynamic changes in global protein sumoylation on BoB challenge over 30 minutes. Longer term impacts were not considered as BoBs inhibit *trans*-splicing and hence lead to multiple alterations in cell physiology. Moreover, we conducted analysis on a cell line with a GFP-tagged CPSF3 protein, and analysed this by Western blotting using both anti-GFP and anti-SUMO antibodies (Figure 4).

**Figure 4:**
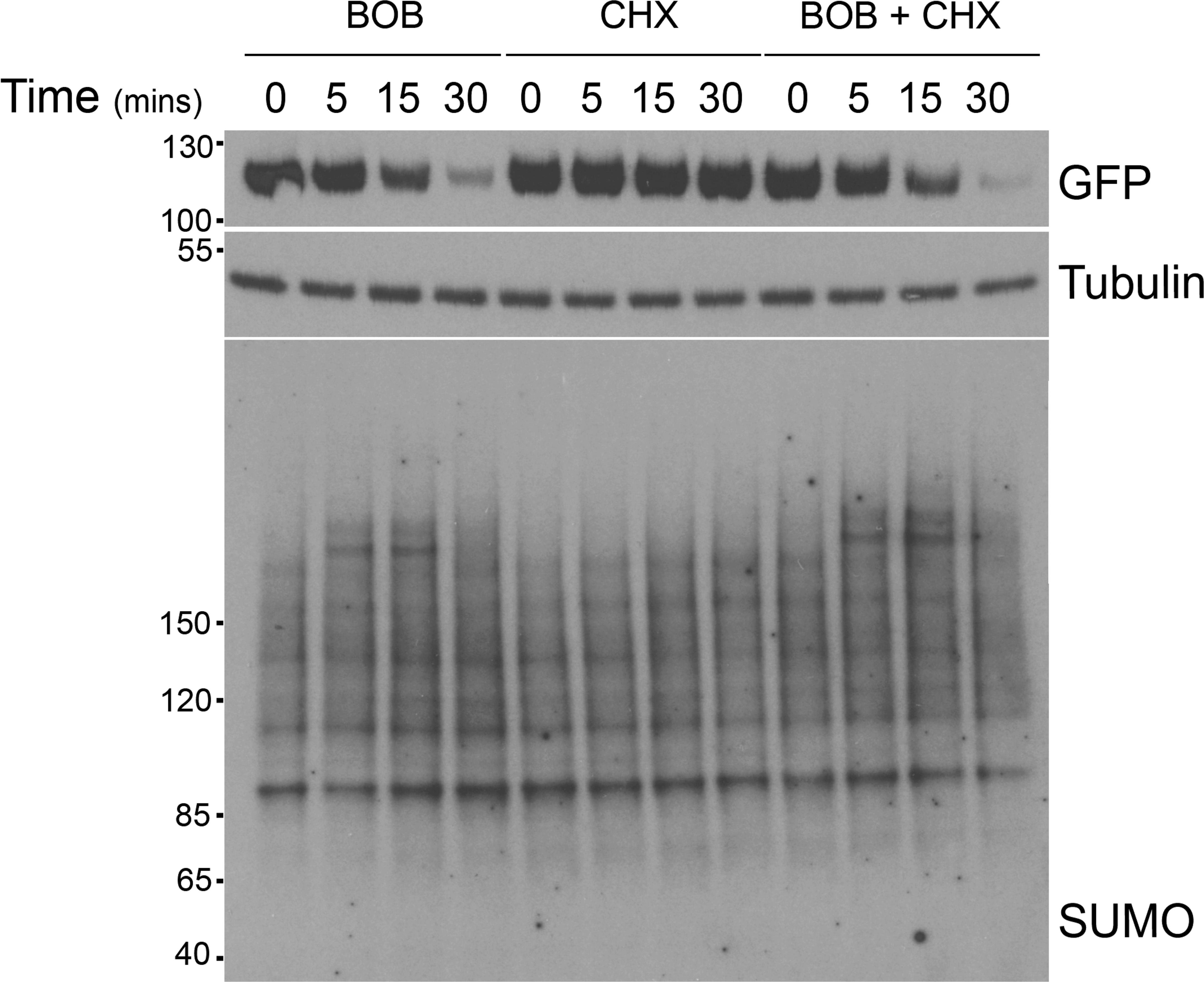
Benzoxaboroles lead to rapid increases in sumoylation and destabliisation of CPSF3. Western analysis of whole cell lysates of cells harbouring CPSF3::GFP and exposed to benzoxaborole (BoB) and/or cyclohexamide (CHX) over a 30 minute period. Top are Western blots for GFP and tubulin, demonstrating rapid degradation of CPSF3 following BoB exposure. The main panel shows an increase to global sumoylation that is only seen with the addition of BoB and not CHX. Migration positions of co-electrophoresed molecular weight markers (in kDa) are shown at left and the antibody used at right.

First, we observed the rapid loss of CPSF3 protein specifically induced by exposure to a benzoxaborole (Figure 4). This was not due to CPSF3 being an inherently unstable protein as, after 30 minutes, there was an insignificant level of degradation when protein synthesis was blocked with cyclohexamide in otherwise untreated cells. Further, cylcohexamide treatment did not protect against enhanced benzoxaborole-mediated CPSF3 turnover. This rapid impact was mirrored in global levels of sumoylation, where it was apparent that an increase was observed as rapidly as after only five minutes. Notably this enhanced sumoylation was only retained for a short period, as at 30 minutes the sumoylation profile had essentially returned to pre-treatment levels. Importantly, a similar impact was not seen with cyclohexamide and therefore this transient hypersumoylation is not the direct result of blocking protein synthesis. Interestingly, this is reminiscent of the sumoylation response to heat shock and oxidative stresses in other eukaryotes, where multisubunit protein complexes become hypersumoylated (Moallem et al., 2023, Shao et al., 2023).

### Destabilisation of CPSF3 is specific to benzoxaboroles and proteasome-dependent

We next asked if multiple benzoxaborole derivatives destabilise CPSF3 and if a second drug, suramin, which elicits a stress response (Zoltner et al., 2020), affects CPSF3 turnover (Figure 5). As expected, multiple benzoxaborole derivatives destabilised CPSF3 within ten minutes. This effect is fully prevented by MG132, indicating that the proteasome is required for turnover (Figure 5A). Moreover, suramin had no impact on levels of CPSF3, indicating that a general stress response is unlikely to be responsible for the enhanced CPSF3 degradation (Figure 5B). To demonstrate that components of the trypanosome sumoylation pathway mediate CPSF3 degradation, we silenced both the SUMO E1 activating and E2 conjugating enzymes (Figure 5B and C). In these SUMO-ligase silenced cells BoB-mediated enhanced CPSF3 turnover was prevented (Figure 5C).

**Figure 5:**
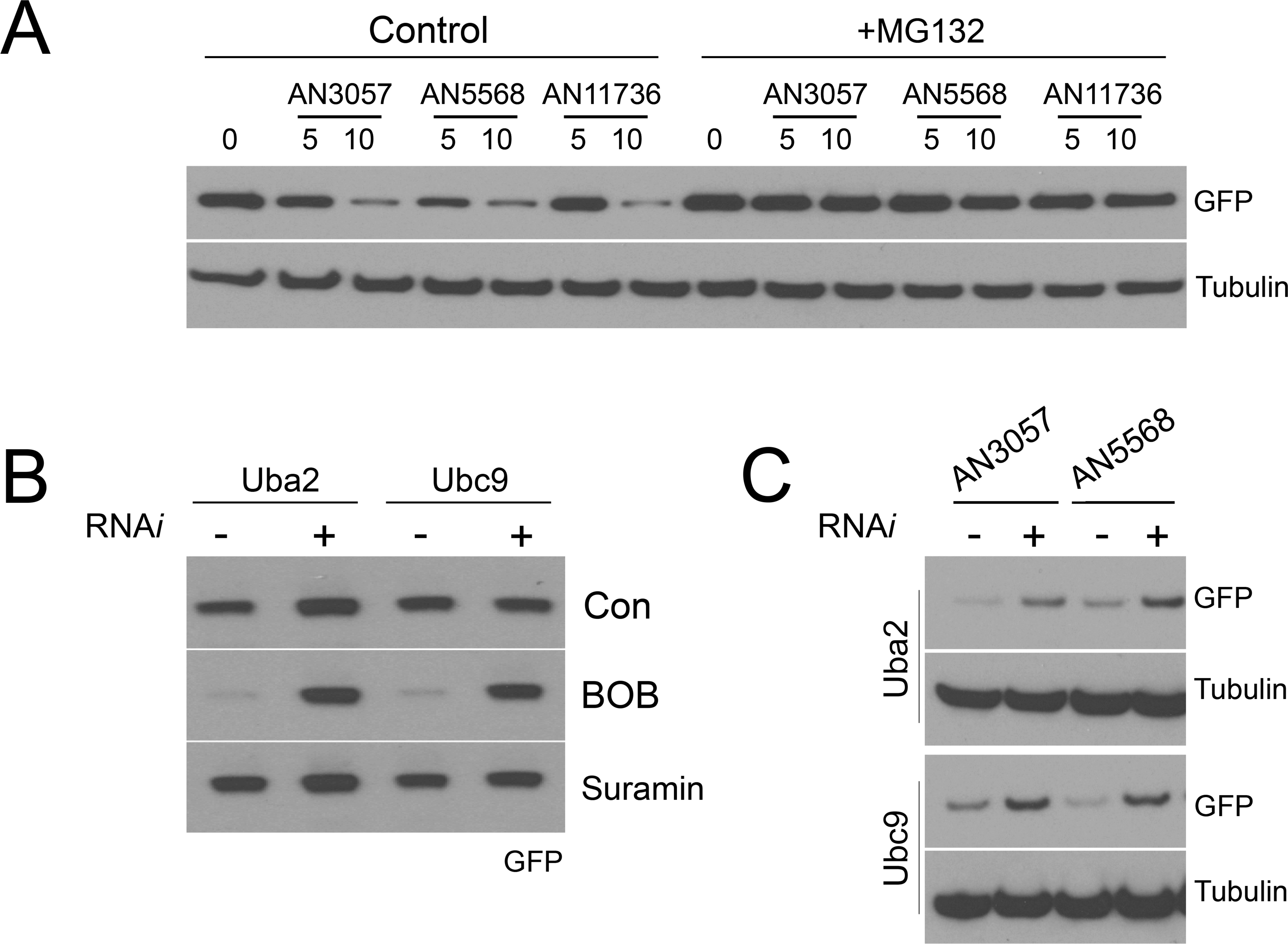
Enhanced degradation of CPSF3 is proteasome and sumoylation-dependant. A: Western blots for CPSF3::GFP in cells treated with benzoxaborole and / or MG132. Cells were exposed for up to ten minutes to BoB and pretreated for 30 minutes with MG132 as indicated and analysed using SDS-PAGE and anti-GFP antibodies. B and C: Turnover of CPSF3 is mediated by sumoylation. Western blots for CPSF3::GFP demonstrating stabilisation of the chimera in cells with SUMO pathway components Uba2 and Ubc9 silenced. As a stress control, cells were also treated with suramin. Blots were probed with GFP and tubulin antibodies as indicated.

### CPSF100, an additional component of the CPSF complex, is also destabilised by benzoxaboroles

CPSF3 functions as the sole catalytic component of the CPSF complex and is crucial for pre-mRNA maturation. As CPSF3 is a direct target of benzoxaborole treatment we asked if additional CPSF complex components were affected. CPSF100 physically interacts with CPSF3 and is conserved in trypanosomatids (Sullivan et al., 2009). Hence, to examine this we endogenously tagged CPSF100 with 3-flag and 6-his epitopes at the C-terminus.

Cells were exposed to three structurally divergent benzoxaboroles and in all three cases the impact was essentially identical. CPSF100 presented as a monomer at ∼100 kDa, together with additional species migrating at >170 kDa (Figure 6), likely a reflection of SDS-resistant oligomers and/or an association with additional CPSF components. There was a clear decrease in the levels of all of these flag-epitope containing species upon benzoxaborole exposure. This indicates that benzoxaboroles have a more profound impact on the CPSF complex beyond CPSF3 destabilisation.

**Figure 6:**
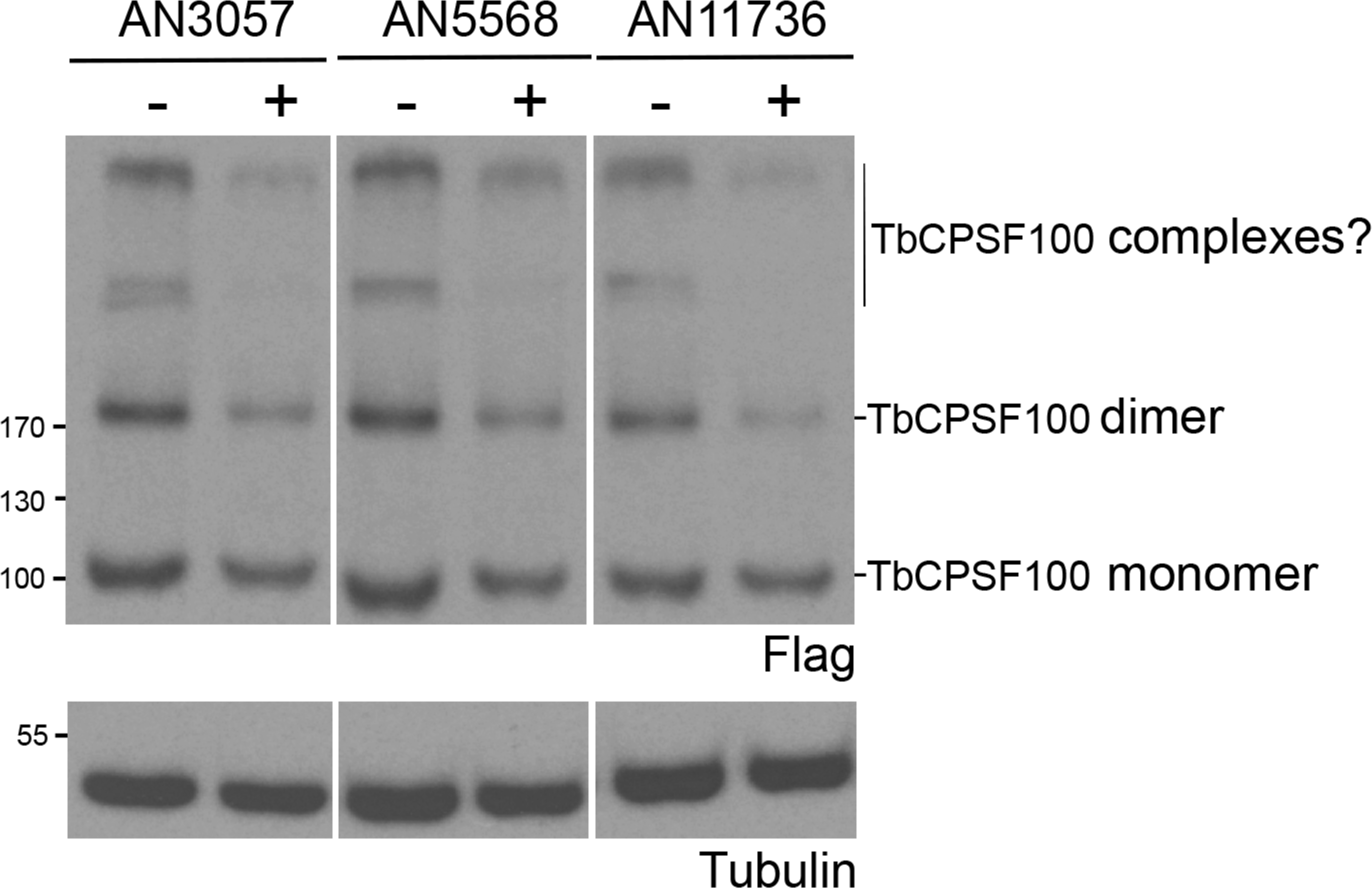
Benzoxaboroles destabilise an additional CPSF complex component. Cells harbouring CPSF100::Flag tagged at the endogenous locus were exposed to benxoxaboroles for 60 minutes and whole cell lysates separated by SDS-PAGE and probed with an anti-flag antibody. Migration positions of co-electrophoresed molecular weight markers (in kDa) are shown at left and assumed migration positions of CPSF100 monomers, dimers and higher molecular weight complexes indicated at right.

### CPSF3 remains nuclear under benzoxaborole treatment

To examine the impact of benzoxaborole on CPSF3 location we examined cells harbouring GFP-tagged CPSF3 and counterstained for SUMO. CPSF3 was present as extensive puncta within the nucleus, while SUMO was observed as diffuse except for a prominent single spot in most nuclei; this latter is the hypersumoylated body (HSB) that locates close to the expression site body (Saura et al., 2014). We observed no obvious relocation of the CPSF3 signal, while the HSB remained prominent but became disrupted at later times (Figure 7A). Moreover, we observed that the distribution of DAPI in 30 minute BoB-treated cells indicated loss of an obvious nucleolus. An apparent increase in the SUMO signal in cells at 30 minutes may be a result of this and increased access of the anti-SUMO antibody.

**Figure 7:**
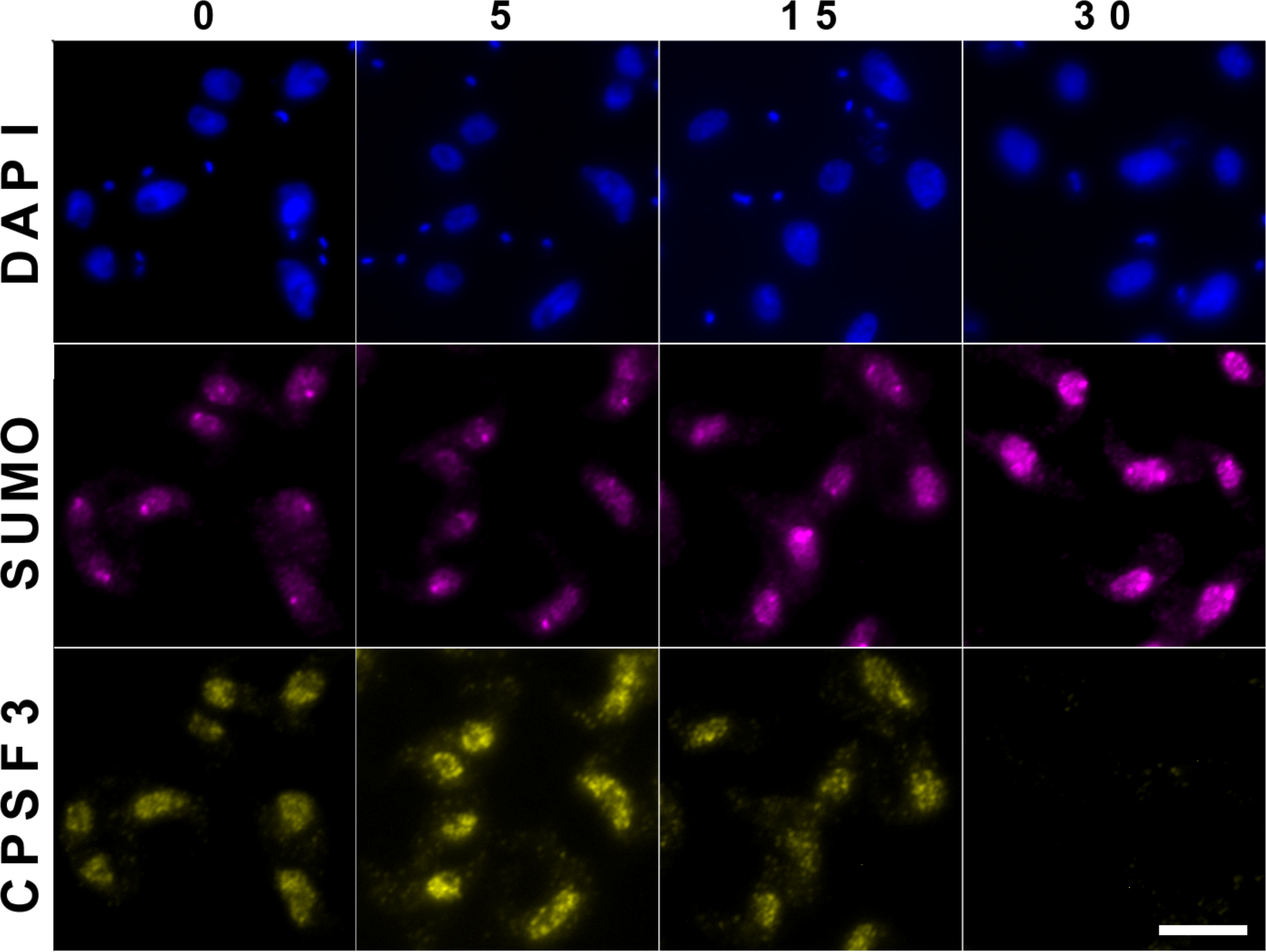
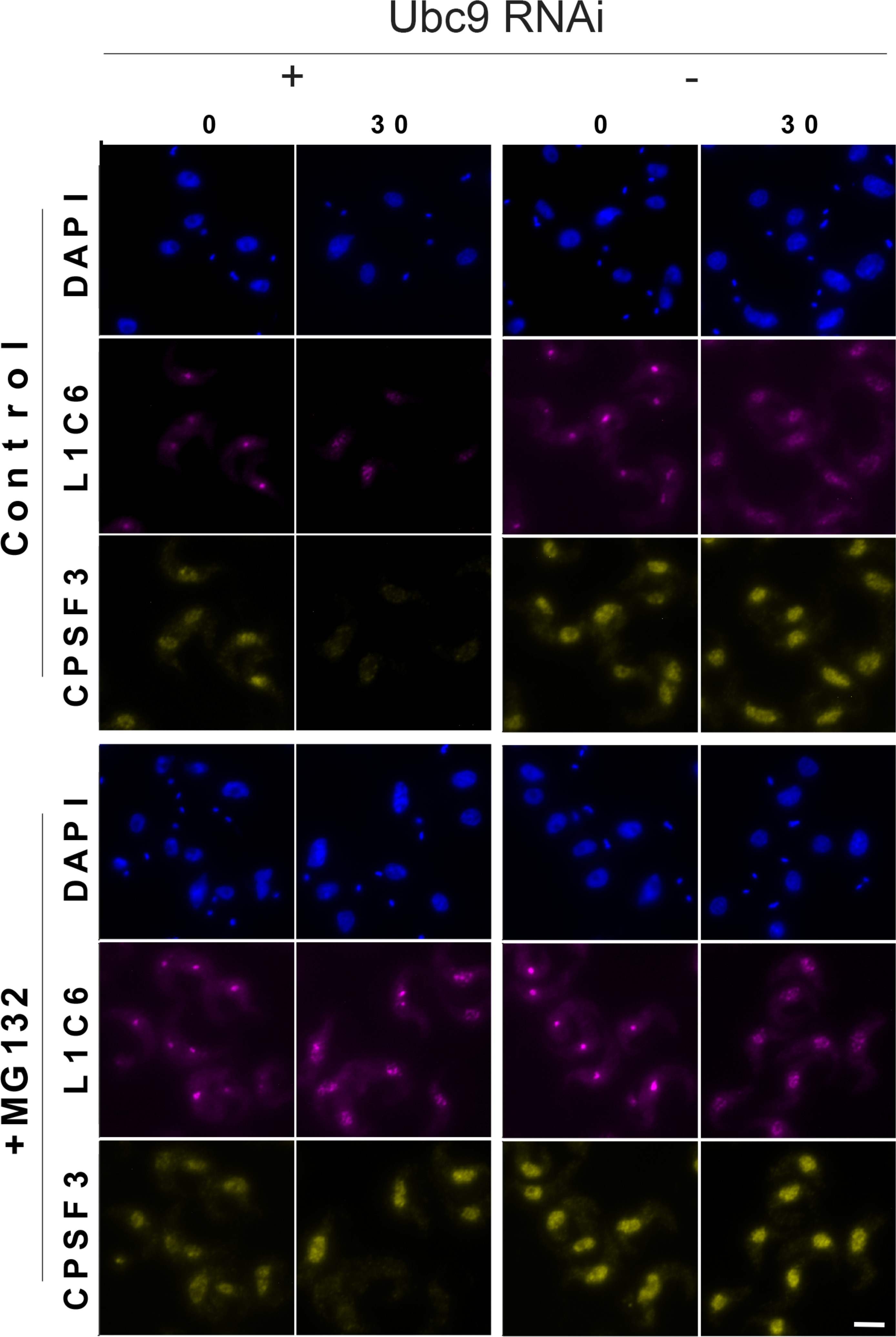
Benzoxaboroles impact both stability and location of CPSF3 and the nucleolus. A: Cells harbouring CPSF3::GFP were treated with benzoxaborole for up to 30 minutes. The cells were counterstained with DAPI for DNA and anti-SUMO antibody. B: Cells harbouring CPSF3::GFP and a Ubc9 silencing construct were treated with benzoxaborole with and without MG132 pretreatment for 30 minutes. The cells were counterstained with DAPI for DNA and the nucleolar marker L1C6 antibody. Scale bar is 2µm.

To determine the impact of the sumoylation pathway on CPSF3 localisation and also to validate apparent loss of the nucleolar compartment we examined cells with a tagged CPSF3 as well as silenced E2. We also counterstained with L1C6 monoclonal, a marker for the trypanosome nucleolus (Durand-Dubief and Bastin 2003). CPSF3 was stabilised by silencing of the E2 enzyme UBC9 and also MG132 exposure, confirming the role of the proteasome in CPSF3 degradation (Figure 7B). Further, the impact of BoB-treatment on the nucleolus was confirmed, but this was not prevented by SUMO E2 silencing. Interestingly, MG132 also led to some fragmentation of the nucleolus when present in combination with BoB, and this was enhanced by E2 silencing. This latter effect may well be secondary as a result of an impact on mRNA processing.

### Benzoxaboroles destabilise multiple CPSF complex components

Given these profound impacts of benzoxaboroles on nuclear organisation, sumoylation and demonstration that stability of more than one component of the CPSF complex is impacted, we analysed global proteome changes in whole cell lysates. To focus on the primary impact of the hypersumoylation response, we undertook liquid chromatography coupled to tandem mass spectrometry (LCMSMS) analysis at 15 minutes post-treatment and compared with untreated cells by stable isotope labelling in cell culture (SILAC) proteomics. We quantified 3,033 proteins, approximately 40% of the total proteome, in which there were 53 proteins with increased and 213 proteins with decreased abundance (>1.2 fold), a remarkable response given the short time of drug treatment (Figure 8, Table 1, Table S3).

**Figure 8:**
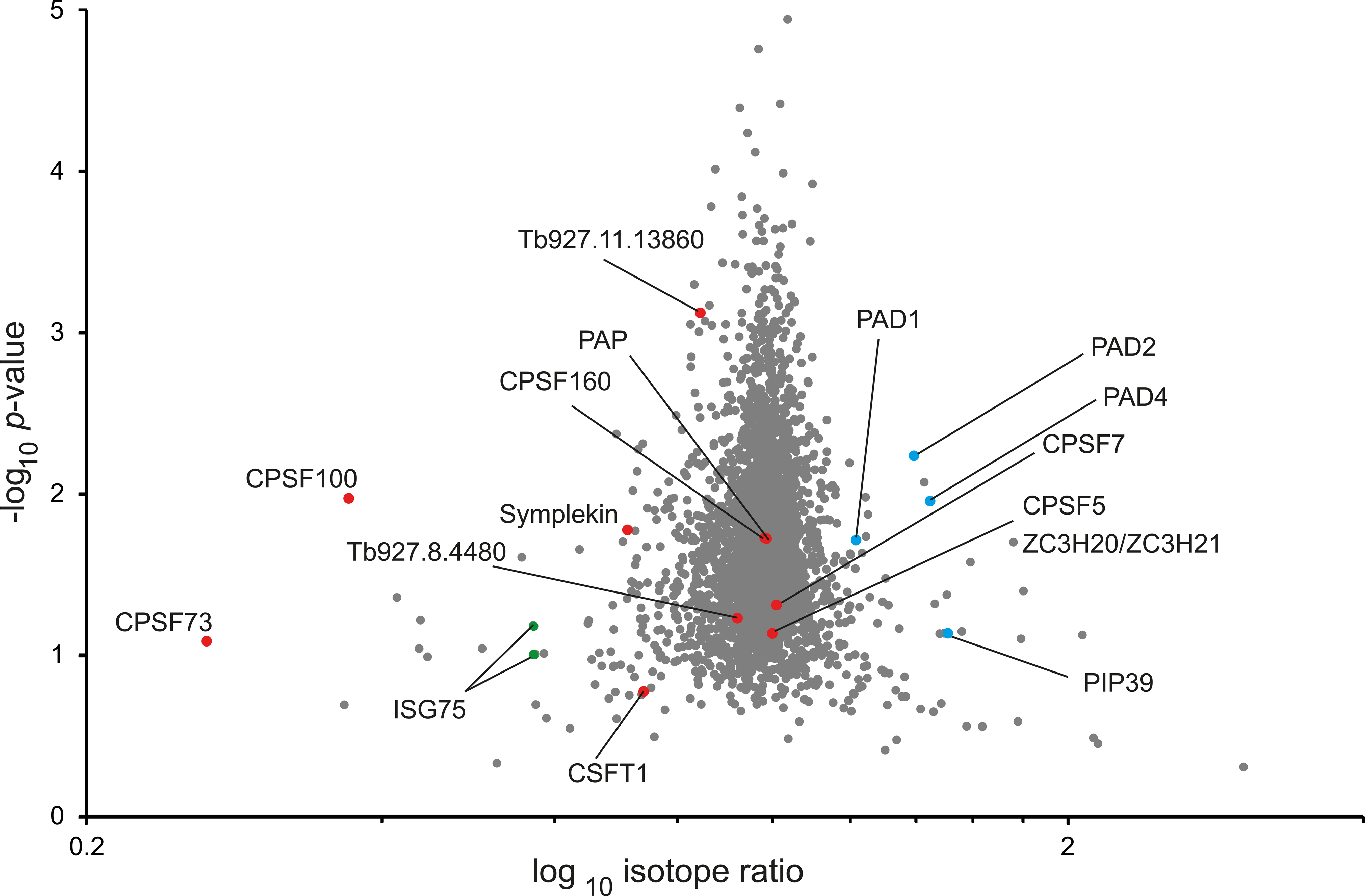
Global changes to proteome highlight splicing components and stress/ differentiation pathways. Cells treated with AN3057 for 15 minutes were harvested and analysed for protein abundance compared to untreated cells. Data points in the volcano plot are coloured red for components of the CPSF complex, green for ISG75 and blue for proteins associated with differentiation/stress responses. Note the coordinated increase in differentiation/stress-associated proteins and the decrease in CPSF complex components. Tb927.8. 4480 and 11.13860 are both nuclear-localised proteins and are restricted to the kinetoplastids.

Four members of the CPSF complex were altered in abundance. Two of these, CPSF3 and CPSF100, are highly significant and confirm earlier Western blot evidence. The two additional CPSF complex components, CSTF1 and Symplekin are clearly decreased but with a lower amplitude. Additional components of the CPSF complex were essentially unaltered in this experiment, suggesting that the CPSF3 subcomplex is hypersensitive to benzoxaborole treatment. Importantly, three of these components, CPSF73, CPSF100 and Symplekin are also components of the histone-specific mRNA processing pathway in mammalian systems (Dominski et al, 2005; Sun et al., 2020); the absence of CSTF1 from this latter complex perhaps points to an impact restricted to the more general mRNA processing CPSF complex, as does the absence of evidence for altered abundance of any potential histone mRNA processing complex-specific factor. An additional kinetoplastid-specific CPSF complex protein, Tb927.11.13860 (Koch et al., 2016), was also slightly decreased. Further, four other CPSF complex proteins were identified, CPSF5/NUDT21, CPSF160, CPSF7/CFIm59 and Tb927.8.4480 (a trypanosome-specific CPSF160-interacting protein) (Koch et al., 2016) but are unchanged in abundance. Tb927.8.4480 and Tb927.11.13860 are of near identical length and have weak homology which may suggest that they are ancient paralogs (Figure S2). Once more, the very short exposure time may explain this lack of impact or that the distinct subcomplexes are differentially regulated.

We also observed increased abundance of differentiation/stress-associated proteins PAD1, PAD2, PAD4 and PIP39. Several of the mRNAs from this cohort interact with RBP10, in common with Tb927.9.15290 that is downregulated, further supporting a connection between the SUMO pathway and differentiation (Mugo and Clayton 2017). Finally, the zinc finger protein ZC3H20, which is indistinguishable from its paralog ZC3H21, has increased abundance. ZC3H20/21 are mRNA binding proteins that stabilise mRNAs associated with differentiation from the bloodstream to the procyclic form, and consistent with the increased abundance of the other differentiation markers (Liu et al., 2020).

### Benzoxaborole treatment leads to a rapid decrease in bulk mRNA

The proteomics data suggest a partial destabilization of the CPSF complex. Given that it is known that CPSF complex factors form subcomplexes that can associate with more than one mRNA processing pathway, we asked what impact BoBs have on global mRNA production. We expected a rapid effect given the block to *trans-*splicing and hence we opted to analyse the transcriptome at 20- and 60-minutes post-treatment with compound. Total RNA was isolated at these two treatment timepoints and an untreated control and poly-A-enriched fractions subjected to sequencing on a DNBseq platform, allowing quantification of 8434 mRNAs and pseudogenic transcripts. Indeed, after 20 minutes of AN3057 treatment, the majority of transcripts (5490) were significantly decreased (FDR=0.01) and 7602 after 60 minutes (Figure S3; Table S4). The cohort of transcripts remaining stable over 60 min showed significant enrichment of gene ontology (GO)-terms including transcripts for ribosomal proteins and histones among several smaller functional category groups (Figure S3; Table S4). Comparison to a recent study on trypanosome mRNA decay (Fadda et al., 2014) revealed that this cohort is dominated by mRNAs with half-lives longer than 1 h (Figure 9). Further, the overall impact of BoB exposure on mRNA levels resembled the consequences of combined inhibition of splicing and transcription elicited by sinefungin and actinomycin D (Fadda et al., 2014). In conclusion, BoB treatment has a global and rapid effect on bulk mRNA levels, with no evidence for selective impact on a specific CPSF subcomplex.

**Figure 9:**
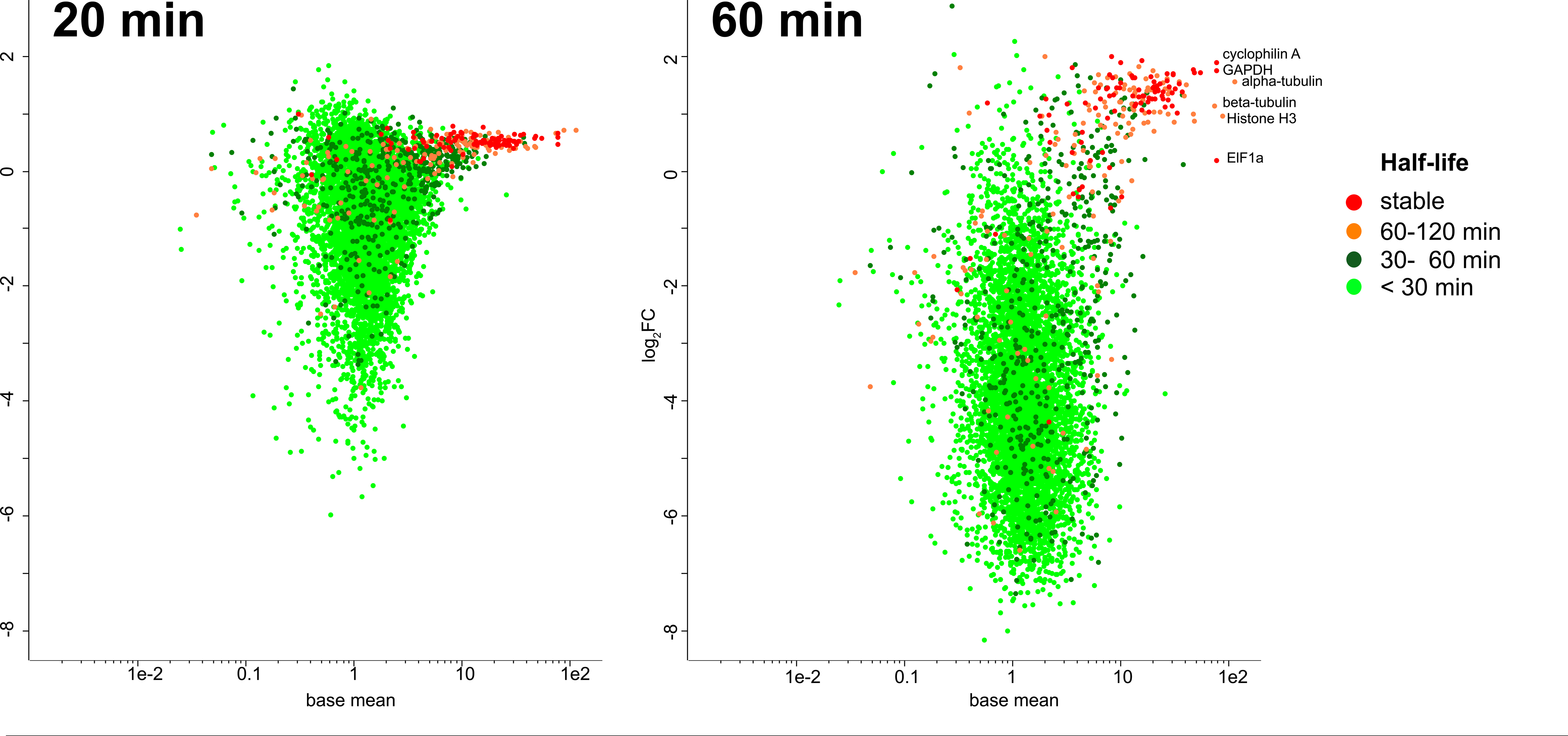
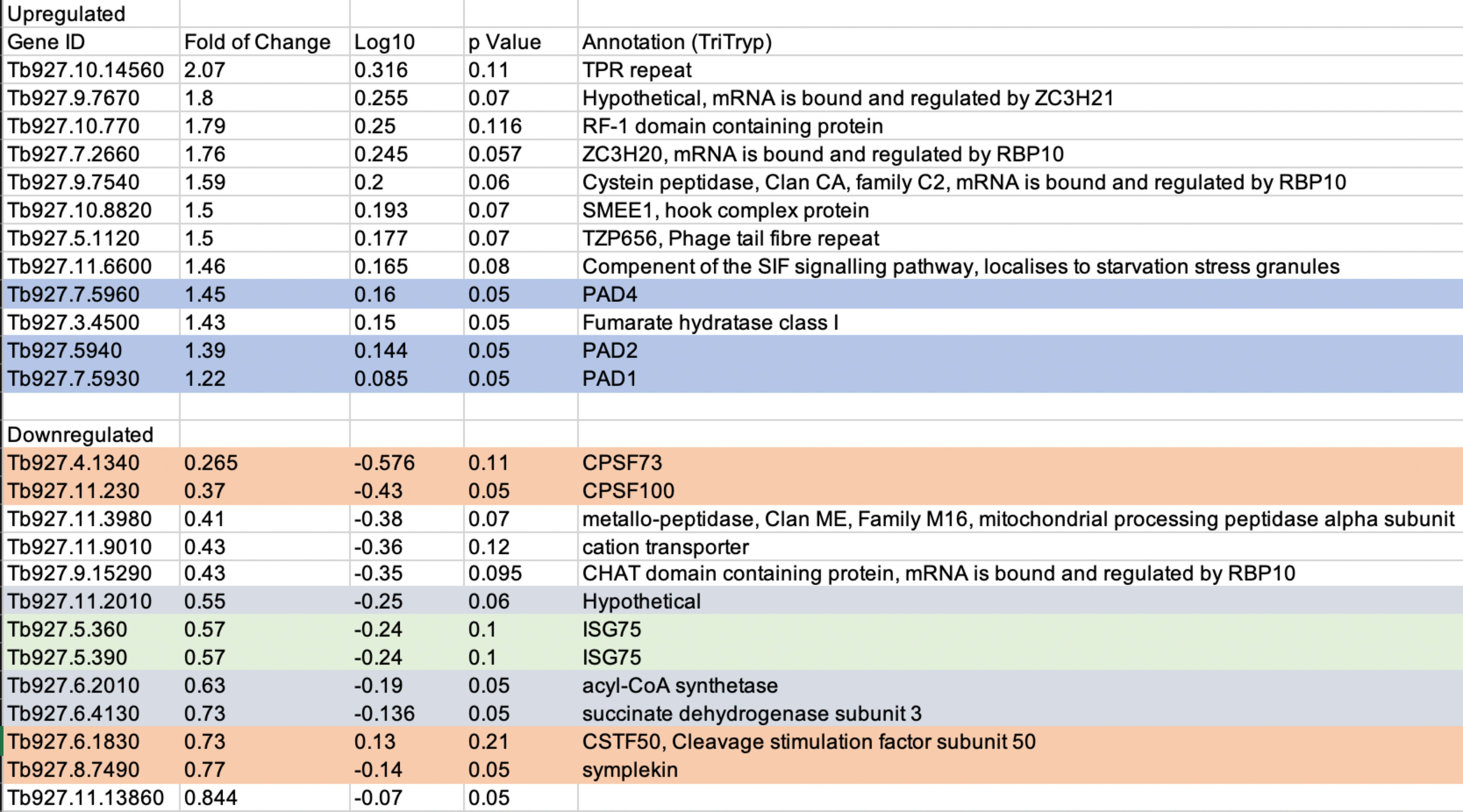
A near global decrease in mRNA levels. The poly-A enriched transcriptomes of cells exposed to AN3057 for 20 minutes and 60 minutes, respectively, were sequenced and compared to untreated control cells (all in 2 biological replicates). For both timepoints, logarithmic fold-changes for 6426 mRNAs are plotted against the length adjusted base mean determined from the control cells. For the mRNAs shown, half-lives were determined previously (Fadda et al., 2014) and those are mapped in color-coded increments (see legend; ‘stable’ denotes transcripts that were not found decayed within 2 h (Fadda et al., 2014)). Selected datapoints are annotated by protein product description. Respective plots including all transcript reads are in Figure S3.

## Discussion

The need for new drugs to challenge infectious disease and other morbidities remains constant and pressing. Benzoxaboroles, with a boron-containing heterocyclic core, offer both a new chemical modality and a number of promising therapeutic avenues; several have entered the clinic (Das et al., 2022). Benzoxaborole indications include the treatment of bacterial, fungal and protozoan infections. Two common targets for infectious agents have emerged, leucyl-tRNA synthetase (LeuRS) and CPSF3, a component of the mRNA maturation system, indicating a common high sensitivity to interference with protein synthesis pathways (Zhang et al., 2019, Sonoiki et al., 2017, Wall et al., 2018). CPSF3 appears to be a common target amongst pathogenic protozoa, and hence understanding BoB interactions with one parasite may well provide broader insights and have implications for other parasites.

Using a combination of biochemical and genetic methodology we demonstrate that, in addition to impact on CPSF3 activity, BoBs also stimulate a sumoylation response, and that this leads to rapid destabilisation of CPSF3, CPSF100 and symplekin, all subunits of the CPSF complex, and essentially inactivating the ability of trypanosomes to process mRNA. Hence BOBs act to target degradation of CPSF3 and hence the overall CPSF complex through stimulating sumoylation, and essentially a targeted protein degradation mechanism. This is fully consistent with the loss of *trans*-splicing activity reported earlier (Begolo et al., 2018), as well as a collapse in protein synthesis. We further demonstrate that BoB treatment has a rapid effect on global mRNA levels, comparable to a global shutdown of transcription and *trans*-splicing induced by combined treatment with actinomycin D and sinefungin (Fadda et al., 2014). A recent study (Sharma et al., 2022) also reported destabilisation of CPSF3 and many other proteins in acoziborole treated trypanosomes, as well as an impact on endocytosis and protein synthesis. However, in that study cells were incubated with BoB for six hours, with the result that many of the observed changes are likely to be secondary impacts arising from the overall blockade to protein synthesis as well as turnover of proteins critical for specific functions, such as endocytosis.

Importantly, we also demonstrate that the impact on sumoylation is dependent only on the presence of the core heterocycle within the BoB, and is relatively insensitive to the presence of additional substituents. Furthermore, degradation of CPSF3 is prevented by silencing of the E1 and E2 SUMO enzymes, indicating that degradation is mediated via sumoylation, as well as through the proteasome. In most organisms, including trypanosomes, sumoylation is a post-translational modification that mainly occurs within the nucleus. In *T. brucei* SUMO is associated with both VSG-encoding chromatin and concentrated close to the active subnuclear region for VSG expression (the ESB). Given spatial proximity between the spliced leader and the active VSG gene in the bloodstream form nucleus, sumoylation is therefore likely a major control factor underpinning antigenic variation. We suggest that there is sufficient conservation of the nuclear pore complex and mRNA processing that it is likely that sumoylation participates both in control of mRNA maturation complexes as well as delivery of RNP complexes to the NPC for their export (Rouviere et al., 2018).

Disruption of the nucleolus also suggests a very rapid impact on ribosomal assembly and possibly rRNA biosynthesis. This is of interest as several mRNAs appear rapidly sensitive to inhibition of protein synthesis, with the suggestion that a short half-life protein is responsible for their degradation (Webb et al., 2005, Dorn et al., 1991). Moreover, as the increase in protein levels for several PAD-proteins and proteins for which RBP10 is a known regulator were notable, this is also similar to the response in *S. cerevisiae* to DNA damage and altered tonicity (Folz et al., 2019), *albeit* that intoxication with suramin does not elicit a sumoylation response. Hence the pathway that is stimulated by BoBs may be related to a differentiation mechanism, or, as we have speculated previously, that general stress responses are able to serve dual duty as differentiation mechanisms (Quintana et al., 2021).

In summary, we demonstrate rapid targeted protein degradation of CPSF3 via sumoylation is associated with BoB treatment of trypanosomes, and that this leads to the rapid collapse of RNA processing and protein synthesis. It is unclear if the primary event here is direct inhibition of CPSF3 or its degradation, but we suggest that this double impact to CPSF complex activity explains the high sensitivity of trypanosomes to these compounds. It will also be of interest to understand if sumoylation pathways are impacted in other contexts of BoB utility and what the consequences of such an interaction have on both the infectious agent and the host.

## Methods and materials

### RIT-seq and data analysis

Genes responsible for drug sensitivity were identified using an RNAi library screen (Alford et al., 2013). A cell population harbouring the library was grown in the presence of tetracycline at 1 mg/ml one day prior to benzoxaborole challenge at ∼5x EC_50_. Cultures were maintained and supplemented with drug before extracting DNA. RNAi target sequences were amplified and the collective products were then subjected to high-throughput sequencing. Sequencing, informatics, to map reads to the genome and validation by individual, targeted RNAi was performed as described (Zhang et al., 2018).

### Drug potency (EC_50_)

EC_50_ was determined by exposing cells to serial dilutions with the highest concentration at 10 µM unless specified otherwise. The assays were conducted in 96-well plate format with three replicates for each sample and a non-treatment control in parallel as described (Zhang et al., 2018). Each EC_50_ curve is representative of three independent experiments (n = 3) and for RNAi two independent cultures analysed.

### Plasmid constructs and cell culture

Bloodstream *T. brucei* (Lister 427) were propagated in T-25 vented cap flasks (Corning Inc., Lowell, MA) at 37°C and 5% CO_50_ with humidity in HMI-9 medium and maintained at a cell density less than 10 cell/ml (Hirumi and Hirumi, 1989). Media was supplemented with 10% foetal bovine serum (v/v) and 1% antibiotic solution (v/v) (Sigma Aldrich). For RNAi a gene-specific target sequence was selected through RNAit and introduced into 2T1 cells using the pRPa^iSL^ plasmid as described (Zhang at al., 2018). All tags used for Western blotting were introduced using either the pPOT or the pNATx^iSL^ vector. The integrity of all constructs was verified by sequencing and all tagged ORFs validated by Western blotting.

### Chemicals, drugs

AN3057 and all other benzoxaboroles (BoBs) were a kind gift from Anacor LLC, or were custom synthesised. Compounds were stored as high concentration stocks in anhydrous MS grade DMSO (Pierce) at -80°C and diluted on the day of use. Cells were treated with BoBs at ∼5x EC_50_, unless stated otherwise, for the times indicated. All other compounds were purchased from Sigma/Aldrich and used at concentrations as described previously (Zhang et al., 2018, Yamada et al., 2023).

### Immunofluorescence

Cells were fixed, permeabilised and nonspecific sites blocked as described (Zhang et al., 2018). Primary antibodies were used at the following dilutions: anti-SUMO (gift from Miguel Navarro, Grenada) at 1:100, anti-mNG (GFP) 1:100 and monoclonal L1C6 (gift from Keith Gull, Oxford) at 1:50. Secondary antibodies (Life Technologies) were Alexa Fluor 568 conjugated anti-rabbit IgG (1:1000) and Alexa Fluor 488 conjugated anti-mouse (1:1000). Coverslips were mounted using *Prolong Gold* supplemented with 4’,6-diamidino-2-phenylindole (DAPI) (Life Technologies). Specimens were examined on a Zeiss Axiovert 200M microscope and images captured with an Axio-Cam MRm camera. Digital Images were captured and processed using Zen Pro software (Zeiss), Omero (https://www.openmicroscopy.org/index.html) and Adobe Photoshop (Adobe Systems Inc.).

### Western blotting

Proteins were separated by electrophoresis on precast gradient SDS-polyacrylamide gels and transferred to polyvinylidene difluoride (PVDF) membranes (Immobilon; Millipore) using a wet transfer tank (Hoefer Instruments). Non-specific binding was blocked with Tris-buffered saline with 0.2% (v/v) Tween-20 (TBST) supplemented with 5% (w/v) freeze-dried milk and antigens detected by Western immunoblotting. PVDF membranes were incubated in primary antibody diluted in TBST with 1% (w/v) freeze-dried milk for 1 hr at room temperature. Detection was by chemiluminescence with luminol (SigmaAldrich) on BioMaxMR film (Kodak). Antibodies were used at the following dilutions: monoclonal anti-HA (sc-7392, Santa Cruz) at 1:10,000, KMX-1 anti-β-tubulin at 1:2000 (Millipore), anti-SUMO at 1:1000, antiFLAG at 1:500 (ThermoFisher) and P4D1 anti-ubiquitin at 1:1000 (Santa Cruz). Commercial secondary anti-rabbit peroxidase-conjugated IgG (A0545 Sigma Aldrich) and anti-mouse peroxidase-conjugated IgG (A9044 Sigma Aldrich) were used both at 1:10 000.

### Metabolic labelling

Cells were pelleted for 10 min at 4°C, washed twice in PBS and resuspended in 500µl of met/cys-free RPMI-1640 medium supplemented with dialysed FBS followed by incubation at 37°C for 1 hr to deplete endogenous met and cys. Cells were labeled at 37°C for 1 hr with EasyTag EXPRESS (PerkinElmer) at a specific activity of 200 µCi/ml and then instantly cooled on ice prior to extraction and fractionation by SDS-PAGE.

### Proteomics

Stable isotype labeling by amino acids in cell culture (SILAC) labeling and downstream processing and LCMSMS was performed as described (Zoltner et al., 2020). HMI-11 for SILAC (Zoltner et al., 2015) contained either normal l-arginine and l-lysine (HMI11-R0K0) or L-arginine U-^13^C_6_ and L-lysine 4,4,5,5-^2^H_4_ (HMI11-R6K4) (Cambridge Isotope Laboratories) at concentrations of 120 and 240 µm, respectively. Cells treated with 400 nm AN3057 (3 × EC_50_ determined after 24h) were grown in parallel with nontreated cells, in the presence of HMI11-R0K0 or HMI11-R6K4, respectively. Cultures in logarithmic growth phase were mixed after 24 and 48 h, respectively, immediately harvested by centrifugation, washed twice with PBS containing Complete Mini Protease Inhibitor Mixture (Roche Applied Science), resuspended in Laemmli buffer containing 1 mm DTT, and stored at −80 °C. Samples were generated in duplicate, and one label swap was performed. Samples were sonicated, and aliquots containing 5 × 10^6^ cells were separated on a NuPAGE bis-tris 4–12% gradient polyacrylamide gel (Invitrogen). The sample lane was divided into eight slices that were excised from the Coomassie-stained gel, destained, and then subjected to tryptic digest and reductive alkylation. Liquid chromatography tandem MS (LC-MS/ MS) was performed by the Proteomic Facility at the University of Dundee on an UltiMate 3000 RSLCnano System (Thermo Fisher Scientific) coupled to a Q Exactive HF Hybrid Quadrupole-Orbitrap (Thermo Fisher Scientific). Mass spectra were analyzed using MaxQuant (Cox and Mann, 2008) searching the *T. brucei brucei* 927 annotated protein database from TriTrypDB (Shanmugasundram et al., 2023). Minimum peptide length was set at seven amino acids, isoleucine and leucine were considered indistinguishable, and false discovery rates of 0.01 were calculated at the levels of peptides, proteins, and modification sites based on the number of hits against the reversed sequence database. SILAC ratios were calculated using only peptides that could be uniquely mapped to a given protein. When the identified peptide sequence set of one protein contained the peptide set of another protein, these two proteins were assigned to the same protein group. *P*-values were calculated, applying *t*-test–based statistics using Perseus (Tyanova et al., 2016). Proteomics data have been deposited to the ProteomeXchange Consortium via the PRIDE partner repository (Perez-Riverol et al., 2019) with the data set identifier PXD049328.

### Transcriptomics

*T. brucei* BSF cells in logarithmic growth phase were treated with 400 nM AN3057 (3 × EC_50_ determined after 24h) and placed in an ice-water mix for 5 min after 20 min and 60 min drug exposure. 6.5 x 10^7^ cells were harvested by centrifugation (1000*g, 4°C) together with nontreated samples prepared in parallel. All samples were in 2 biological replicates. RNA was extracted with the RNeasy RNA extraction kit (Qiagen) following the manufacturer’s protocol. Total RNA yields were 12-16 μg with a RNA quality number (RNQ) > 6 (for quality control data see Figure S. RNA samples were stored at −80 °C and sent to BGI for library preparation and sequencing. After oligo dT enrichment, mRNA was fragmented and subjected to first strand cDNA synthesis using random primers. Second strand cDNA was synthesized with dUTP instead of dTTP. The synthesized cDNA was subjected to end-repair and 3’ adenylation before adaptor ligation. Upon digestion of the U-labeled second-strand template with uracil-DNA-glycosylase the library was PCR amplified. The circularized library was amplified to generate the DNA nanoball (DNB) and sequenced on a DNBSEQ platform. Raw data was filtered using SOAPnuke software (Chen et al., 2018) with filter parameters: -n 0.001 -l 20 -q 0.4 --adaMR 0.25 --polyX 50 -- minReadLen 150. For analyzing the data Linux based soft wares were employed. Index files were prepared with Burrows-Wheeler Aligner (BWA) (Robinson et al., 2011), according to *T. brucei* 927 genome FASTA file in TriTrypDB (release 65) (Shanmugasundram et al., 2023). Subsequently, the sequence was aligned with the respective reference file in BWA. Next, Samtools (Li et al., 2009) was utilized for sorting and indexing the output file. Finally, the total reads were extracted with featureCounts (Liao et al., 2011). DESeq2 (Love et al., 201) from the R package was applied to normalize the reads according to control. Statistical and functional category enrichment analyses were performed using Perseus (Tyanova et al., 2016). Transcriptomics data have been deposited to ArrayExpress (Parkinson et al., 2007) with the accession E-MTAB-13843.

## Supporting information

SuppData Table 1

SuppData Table 2

## Acknowledgements

This work was supported by grants from the Wellcome Trust (204697/Z/16/Z to MCF) and the Medical Research Council (MR/P009018/1 to MCF). We are indebted to the FingerPrints proteomics facility at the University of Dundee for excellent mass spectrometry.

## Supplementary data for

**Responses to benzoxaboroles are mediated by SUMOylation in *Trypanosoma brucei***

### Supplementary figure and table legends

**Figure S1:**
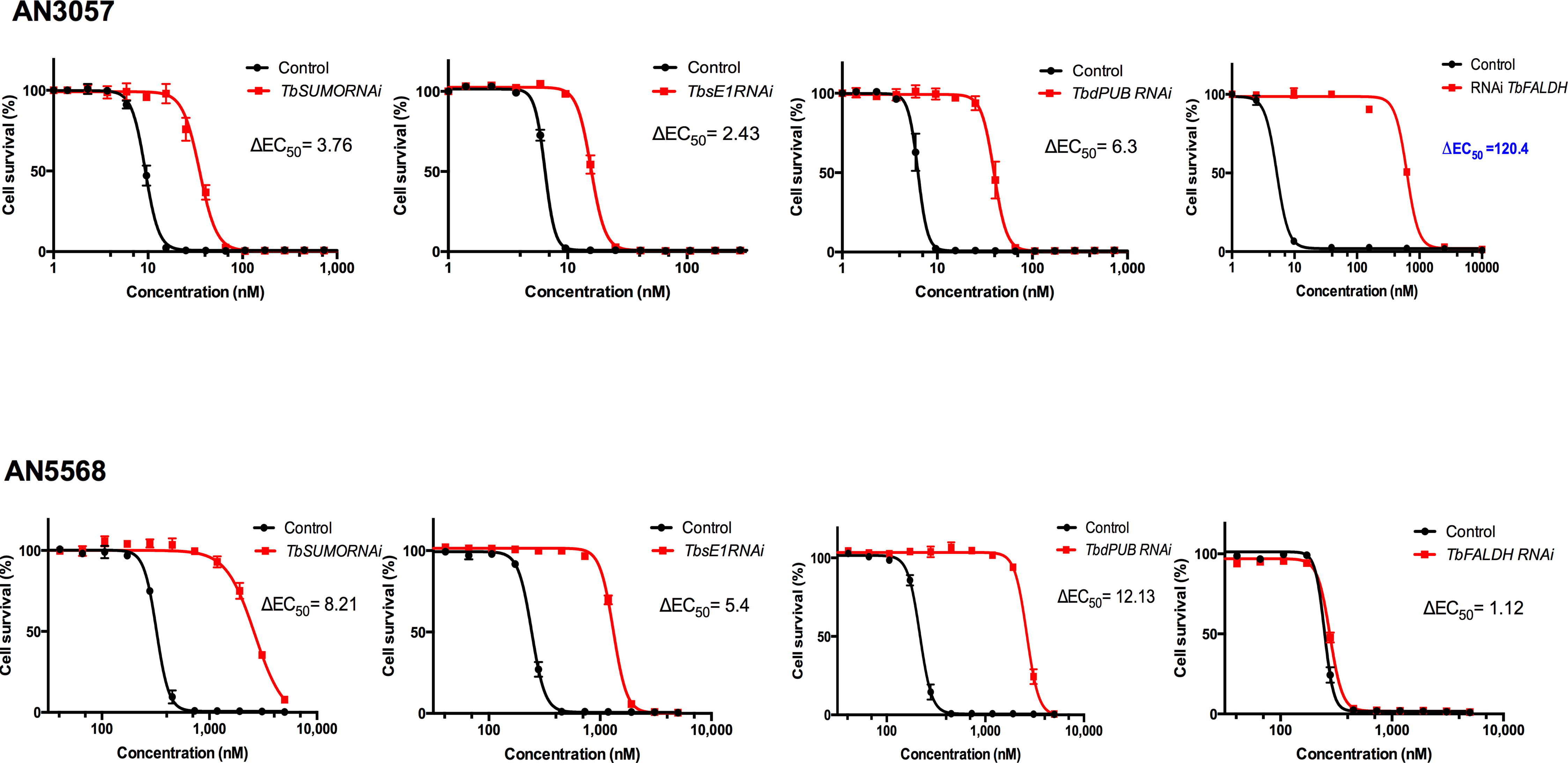
EC_50_ determination for sensitivity towards benzoxaboroles and sumoylation components. Cells were exposed to a range of concentrations of AN3057 and AN5568 in the presence or absence of silencing of sumoylation pathway components. Black curves are control cells and red curves silenced cells.

**Figure S2:**
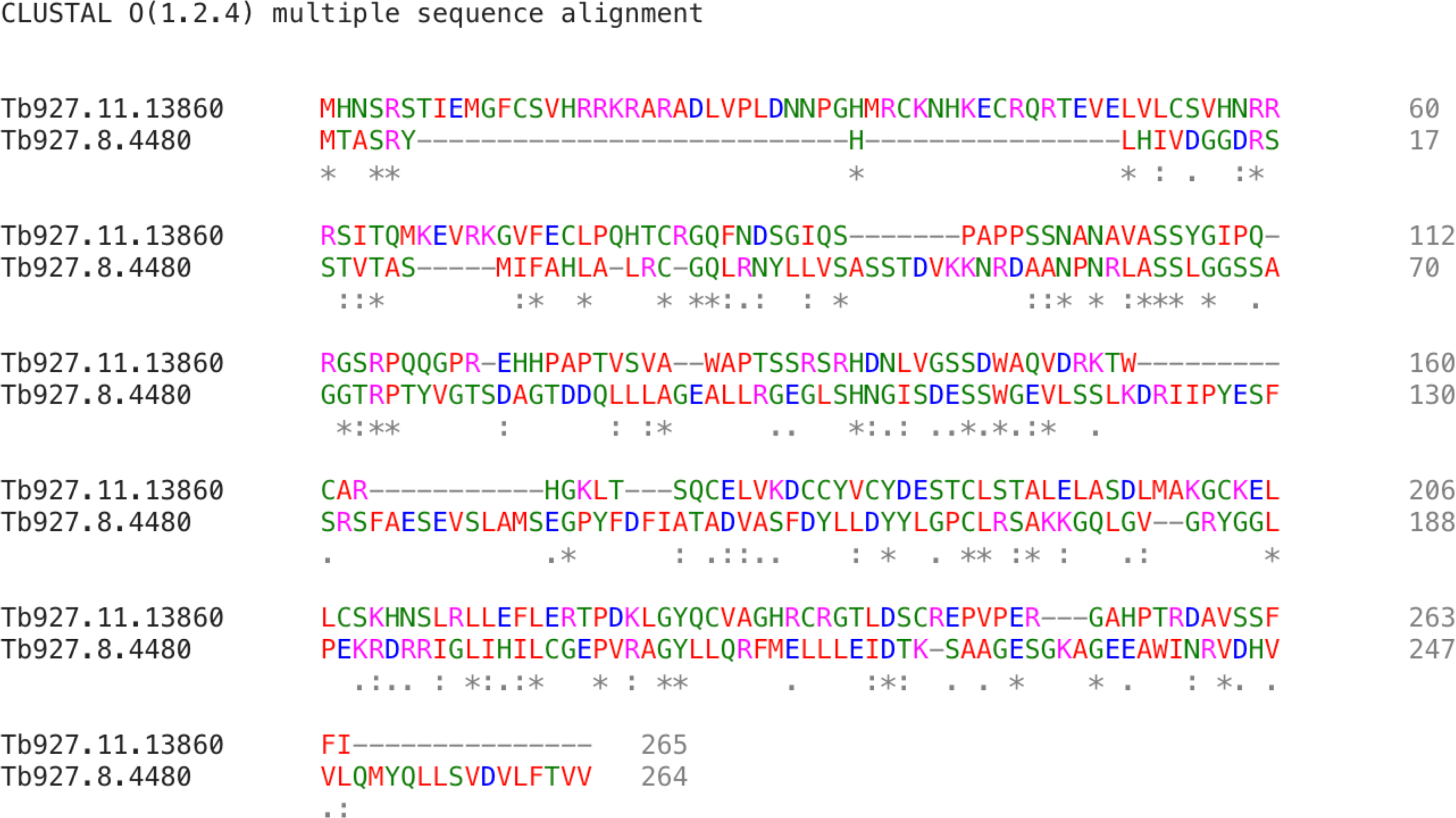
Alignment of predicted protein sequences of Tb927.8.4480 and Tb927.11.13860. Sequences were aligned using Clustal omega and coloured according to residue property using the Clustal convention (https://www.ebi.ac.uk/Tools/msa/ clustalo/).

**Figure S3:**
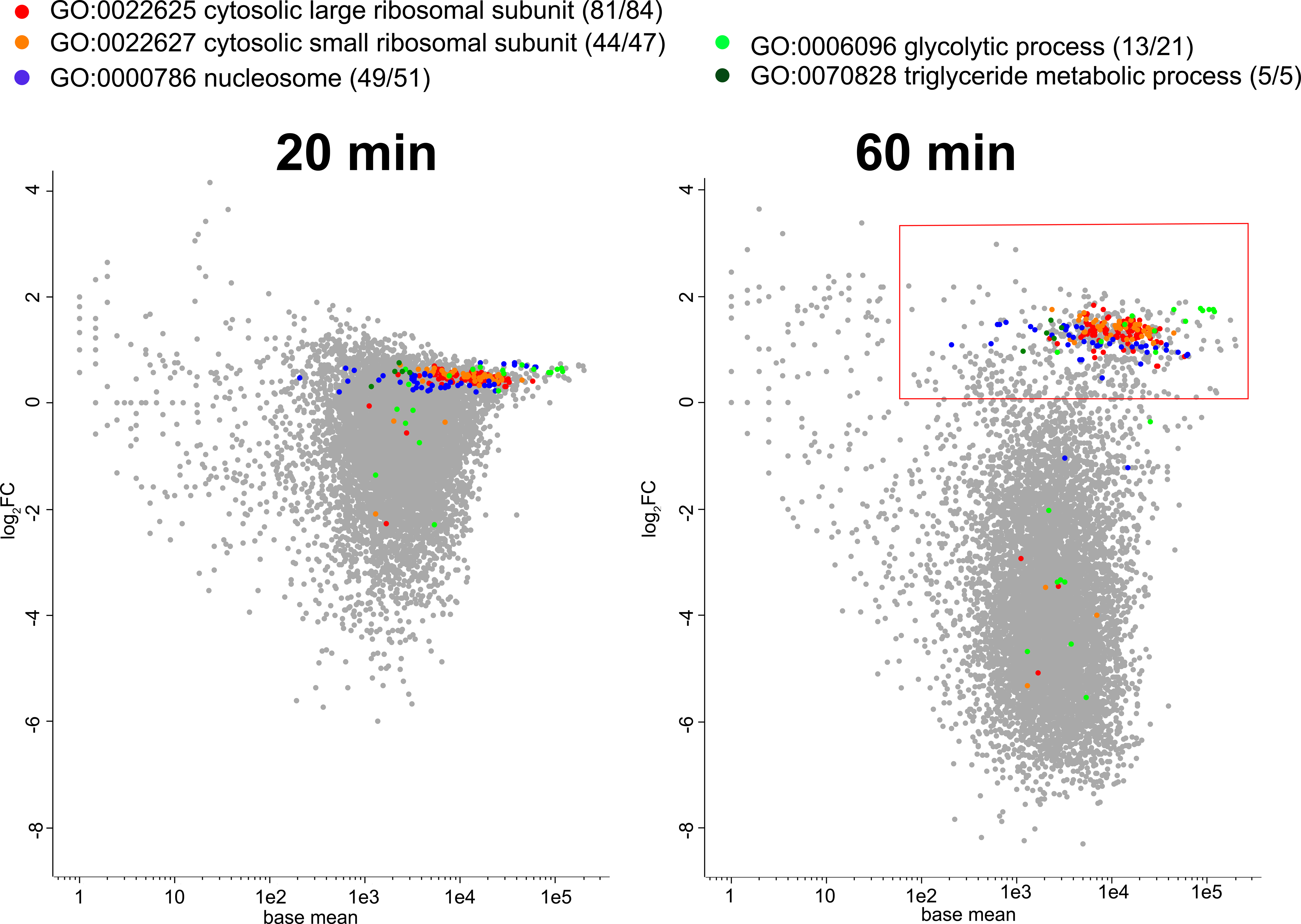
A near global decrease in transcription. The poly-A enriched transcriptomes of cells exposed to AN3057 for 20 minutes and 60 minutes, respectively, were sequenced and compared to untreated control cells (all in 2 biological replicates). For both timepoints, logarithmic fold-changes for 8434 transcripts are plotted against the base mean determined from the control cells. Enriched GO terms for a cohort of 680 transcripts that do not decrease after 60 min treatment (red frame), are indicated by colors for selected curated GO processes; for the full GO term enrichment analysis see Table S3.

**Figure S4:** RNA and transcriptome quality control data.

**Tables S1 and S2. Data for RITseq analysis.** Reads, raw analysis and details for Manhattan plots are provided in these tables and suitable for repurposing.

**Tables S3. Data for transcriptomics analysis.**

**Tables S4. Data for proteomics analysis.**

## Notes

### Competing Interest Statement

The authors have declared no competing interest.

